# Investigating the dynamics of heat acclimation in pig through transcriptome analysis of blood samples

**DOI:** 10.64898/2026.04.01.715954

**Authors:** Guilhem Huau, Laurence Liaubet, Yann Labrune, Paulo Henrique Reis Furtado Campos, Hélène Gilbert, David Renaudeau

## Abstract

This study aimed to investigate the dynamics of gene expression in pigs during heat stress (HS), focusing on both short-term (STHA) and long-term (LTHA) heat acclimation phases. A total of 12 castrated males were exposed to thermoneutral temperatures (24°C) for 14 days (TN) and then to a constant temperature of 30°C for 21 days.

Rectal temperature measurements indicated a biphasic thermoregulatory response, with an initial peak followed by acclimation. Using whole blood transcriptome analysis at seven time points between day 5 before the initiation of HS challenge and day 13 post HS. A total of 525 genes were differentially expressed during the STHA (day 0-day 2) phase. A switch in the expression of most genes was observed around 20 hours after HS. Functional pathway enrichment analysis identified through shape-based clustering revealed the activation of the immune system, especially mediated through toll-like receptor signaling pathways. The LTHA phase (day 2-day 13) revealed 985 differentially expressed genes, with pathways associated with various metabolisms, including mitochondrial fatty acid beta-oxidation, and electron transport, ATP synthesis, and heat production by uncoupling proteins. Interestingly, oxidative phosphorylation was predicted to be activated during the LTHA, particularly in Complex V, whereas other complexes showed mixed regulation. Comparative pathway analysis indicated distinct metabolic adaptations between STHA and LTHA, with up-regulation of glucose and lipid metabolism in late STHA and down-regulation of lipid metabolism during LTHA.

This study contributes to a better understanding of the time course of adaptation mechanisms in pigs to HS, underlying a coordinated regulation during STHA involving several stress-specific mechanisms (via the HSP) and metabolic variation to help pigs achieve homeothermy.

## Introduction

Heat stress (HS) has a significant economic impact on swine producers, especially in hot and humid regions **[1]**. In fact, the economic cost of HS in swine extends beyond direct production losses to include indirect expenses related to the adaptation strategies. The effects of HS are accentuated in modern genetic lines as genetic selection for improving prolificacy in reproductive sows and lean deposition in growing pigs has led to an increase in metabolic heat production **[2]**. According to the projections of the IPCC (IPCC 2023), global warming, as well as an increase in the intensity and frequency of heat waves in key regions for swine production, will increase HS concerns even more in the near future. Heat acclimation (HA) refers to the physiological, biochemical, and molecular adaptations that occur in response to repeated or prolonged exposure to hot temperatures. These adaptations enhance an individual’s ability to tolerate HS and maintain homeostasis. In pigs, HA typically occurs in two distinct phases: an initial period of severe hyperthermia followed by a gradual reduction in internal temperature **[3,4]**. Conceptually, the HA process in homeotherms is described as a transition from an early, transient ’inefficient’ acclimation phase—referred to as short-term heat acclimation (STHA)—to a more ’efficient’ state known as long-term heat acclimation (LTHA) once acclimatory homeostasis has been achieved **[5]**. The acute phase is primarily regulated by homeostatic mechanisms involving the nervous and endocrine systems, while the chronic phase is governed largely by homeorhetic regulators within the endocrine system **[6]**. In pigs, the duration of the STHA and LTHA phases is influenced by the intensity of thermal stress, feeding management, and animal-specific factors, including variability both between and within breeds **[3,7]**. Evidence from rodent studies suggests that the development of the heat-acclimated phenotype during STHA involves the reprogramming of genes responsible for constitutive proteins and stress-inducible molecules **[8]**. To date, and in contrast to other species, the dynamic of gene expression during HS exposure has not yet been studied in swine.

The objective of the present study is to understand the timeline of HS adaptation at a transcriptomic level. Our study focused on monitoring gene expression in blood cells to provide insights into the acclimation responses of the whole body exposed to HS. This study contributes to a larger initiative dealing with identifying blood biomarkers that can indicate adaptive responses to HS in swine production. These biomarkers are key levers for developing effective management strategies to support swine welfare and productivity in hot climates or during periods of elevated environmental temperatures.

## Methods

### Animals and Treatments

This study involved a total of 36 castrated male growing pigs from two divergent lines (n = 19 LRFI and 17 HRFI pigs, respectively) selected for their low (LRFI) and high residual feed intake (HRFI), and was carried out in two consecutive replicates of 17 and 19 pigs, respectively. Residual feed intake is defined as the difference between a pig’s observed and expected feed intake and is a proxy of feed efficiency. The animals used in this experiment were owned by the INRAE (National Research Institute for Agriculture, Food and Environment). These two lines have been selected since 2000 at the INRAE, France, with a process described by Gilbert et al. **[9]**. Pigs from the seventh generation of selection were used in this study. These pigs were raised in selection facilities (INRAE-GenESI **[10]**, Le Magneraud, France), then transferred at weaning to the experimental facilities (INRAE UE 3P **[11]**, Saint Gilles, France). All procedures were conducted in accordance with veterinary best practices, following the AVMA Guidelines for the Euthanasia of Animals (2020) and the ARRIVE guidelines **[12]**. Details of the anesthesia and the euthanasia performed at the end of the study are described in the Surgery section below.

At approximately 80 days of age (i.e., 40 kg of BW), pigs were moved into a temperature-controlled room and housed in individual metal-slatted pens (0.70×2.30 m). Each pen (18 in total) was equipped with a feed dispenser and a nipple drinker to allow free access to water and feed. Diet was formulated to meet the nutritional requirements of this animal category according to standard recommendations **[13]**. Throughout the adaptation period, the room ambient temperature was maintained at 24°C for 14 days. At d_0 of the experiment, the temperature was raised from 24°C to 30°C at a rate of 2 °C/hour starting at 10:00. Therefore, the ambient temperature was maintained for 21 days at 30°C. Photoperiod was fixed to 12 hours of artificial light (08:00–20:00 hours). More details on the experimental design can be found in Campos et al **[13]** and described in S1 Fig.

### Surgery

From the 36 pigs, we randomly selected four and six pigs/line in the first and second replicate respectively for the surgical procedure. These pigs were prepared with a jugular catheter following the protocol previously described by Melchior et al.**[14]**. Two weeks before the start of the experiment, an indwelling silicone catheter (id 1.02 mm, od 2.16 mm; VWR International S.A.S, Fontenay-sous-Bois, France; catalog ref. 228-0656) was implanted through a collateral vein in the right external jugular vein. Sedation was performed using intramuscular ketamine (15-20 mg/kg) and general anesthesia with an oxygen/isoflurane gas mix. A surgical operation was performed under 20 min for each pig. The catheter was placed under the skin up to the back and stored in a strengthened purse sewn on the back of the pig when it was not used for the blood samplings. The catheters were flushed every 2 or 3 days with 5 mL of saline solution containing 5 % of heparin. At the end of the study, all the animals were sacrificed according to protocol by overdose anesthesia with pre-sedation using the following protocol. An initial intramuscular injection of tiletamine/zolazepam at 6-8 mg/kg was given with an 18G x 1.5’’ needle for pre-anesthesia, followed by a slow intravenous injection of embutramide, mebezonium diiodide and tetracaine hydrochloride at 6 to 10 mL depending on live weight.

### Performance and Response Variables Measurements

The rectal temperature (RT) was measured using a digital thermometer (Microlife Corp., Paris, France) in not restrained animals and with the minimum of stress on d-6, d-4, d-1, d1, d2, d3, d6, d8, d10, d15 at 08:30 and 15:00, and on d0 at 09:00, 10:00, 11:00, 12:00, 13:00 and 14:00. To achieve that, during the adaptation period, animals were adapted to the presence of the experimenter in the pen, to the instruments and measurements conditions to avoid any influence of the procedures and measurements by themselves in the observations. From the 20 catheterized pigs, we selected the 12 (6 pigs in each line) having the closest live body weight to the mean of the group. Total whole blood samples were collected at d-5, d0 at 09:00, d0 at 13:00, d0 at 18:00, d1, d2 and d13, between 10:30 and 11:00 hours when not specified, via a jugular catheter after 2 hours of fasting for a better standardization of the blood sampling. Before each sampling, the catheters were flushed with normal saline solution. Afterwards, 2.5 mL of blood were collected and put in PAXgene® (2.5 mL) Blood RNA tubes (PreAnalytiX, QIAGEN/BD, Hombrechtikon, Switzerland). The tubes were then stored at -20 °C.

### RNA Isolation and Microarray Hybridization

#### RNA isolation

Whole blood RNA samples were collected on PAXgene Blood RNA Tubes (PreAnalytiX) and were conserved at -20°C until extraction. Total RNA was extracted according to the manufacturer’s recommendations (PAXgene Blood RNA Kit, Qiagen), and the extracted total RNA was eluted in 40 µl of buffer BR5 and stored at -80°C. Quantification was performed using a Nanodrop to determine the concentration of each purification and the 260/230, 260/280 ratios. Quality was checked by electrophoresis on 0.8% agarose gel. RNA Integrity Numbers (RINs) were determined using a 2100 Bioanalyzer Instrument (Agilent).

#### Microarray data

A specific microarray (8 x 60 K, GPL16524, Agilent Technology) was used, which was a combination of 43,603 probes derived from the 44 K Agilent-026440 porcine-specific microarray (V2, for 71 % of total probes), 3,773 probes from the immune system, 9,532 probes from adipose tissue and 3,768 probes from muscle tissue, as already reported [15]. The microarray probes annotation was updated with the last genome assembly Sscrofa11.1 (nblast/Ensembl 2018) performed by the Sigenae platform (INRAE, Toulouse, France). The annotation of some differentially expressed probes (DEPs) was verified manually by sequence alignment using BLAST against the NCBI or Ensembl databases (version 111).

Only samples with an RNA integrity Number (RIN) > 7 were kept, resulting in 77 samples. The absence of 3 samples from d0 at 09:30 is explained by the poor quality of the extracted RNA (RIN <7). On day 13, failures from some catheters (i.e., clogged catheters) explain the absence of 5 samples. Microarray hybridizations were realized on the INRAE genomic @BRIDGE platform (https://abridge.inrae.fr/, UMR 1313 GABI, Jouy-en-Josas, France). For each sample, Cyanine-3 (Cy3) labeled cRNA was prepared from 100 ng of total RNA using the One-Color Quick Amp Labeling kit (Agilent) according to the manufacturer’s instructions, followed by RNeasy purification (QIAGEN). Dye incorporation and cRNA yield were checked with the Biospec nano spectrophotometer (Shimadzu). 600 ng of Cy3-labelled cRNA (specific activity >8 pmol Cy3/µg cRNA) was fragmented at 60°C for 30 minutes in a reaction volume of 25 µl containing 10x Agilent fragmentation buffer and 25X Agilent blocking agent following the manufacturer’s instructions. On completion of the fragmentation reaction, 25 µl of 2X Agilent hybridization buffer was added to the fragmentation mixture and hybridized to Agilent microarray (8X60K, GPL16524) enclosed in Agilent SureHyb-enabled hybridization chambers for 17 hours at 65°C in a rotating Agilent hybridization oven. After hybridization, microarrays were washed sequentially in Wash buffer 1 (Agilent Technologies, 1 min), Wash buffer 2 (Agilent Technologies, 37°C, 1 min). Slides were scanned immediately after washing on an Agilent G2505C Microarray Scanner with Agilent Scan Control A.8.5.1 software.

The scanned images were analyzed with Feature Extraction Software 10.10.1.1 (Agilent) using default parameters (protocol GE1_1010_Sep10 and Grid: 037880_D_F_2012). All subsequent data analyses were done under R (www.r-project.org) using packages of Bioconductor (www.bioconductor.org). Raw data (median of pixels intensity) were imported into R using the read.maimages function from the limma package. Probes were filtered and kept if they had a good signal in more than 75% of the samples in at least one experimental group. Using this filter, 32,924 probes were selected from a total of 61,625, or 53.4%, normalized by the median, and log2 transformed. Transcriptome data was corrected for the technical batch effect of the chip (each corresponding to 8 microarrays) using the ComBat procedure from the *sva* R package [16]. The resulting matrix has 32,924 rows, each corresponding to a unique ProbeName (provided as data Matrix) and 75 columns (pigs x day +/- hour). Normalized expression data (log2 transformed), raw data, and samples information are available with the accession ID GSE271309 on NCBI/GEO (https://www.ncbi.nlm.nih.gov/geo/).

### Data and Statistical Analyses

Data were analyzed using the R 4.2.2 software [17] in a renv [18] environment with packages from Bioconductor and CRAN to ensure the reproducibility of the analyses.

The RT was calculated as a mean per day using morning (at 08:30) and evening (at 15:00) measures. Then, RT data were analyzed using a two-way mixed ANOVA with the rstatix package [19]. Hypothesis for the ANOVA were tested (normality of the data with a qqplot, homogeneity of variance and covariance using Levene’s and Box’s M-test, respectively). The RFI line was used as the between-subjects variable, and the day of the experiment as the within-subjects variable.

Based on a preliminary analysis, pre-HS data (d-5 and d0 at 09:00) have been pooled in a single value for each probe using the mean of the two pre-HS values. For some animals, only one of the two values (d-5 or d0 at 09:00) was present and was used as reference value. For validating this pooling procedure, a paired t-test followed by a p-value correction for multiple testing (FDR) was used to assess that the majority of the probes have a similar expression during the preHS period.

The differential analysis was performed in two steps to study the expression of genes involved in the short-term HA (d0 vs d2, STHA) and long-term HA (d2 vs d13, LTHA).

In a first step, data was filtered to keep pre-HS and HS data up to 2 days of exposure to assess the acute responses to HS. The dynamic probe expression was modeled (1) with a generalized additive mixed model [20]. For each probe, two spline functions, one for each line, were used to model the expression of the corresponding transcript. The model (1) was defined with the two splines as a function of the hours since the start of the HS, a parametric term for the line according to the recommendation of Wood [21], and the replicate and pig as a random effect.

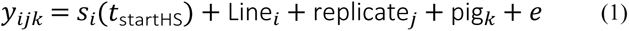

To select the correct hyperparameters, we performed a grid-search for the spline basis (between thin plate regression splines, thin plate regression splines with shrinkage, cubic regression spline, and cubic regression spline with shrinkage from mgcv) the spline basis size, and the smoothing parameter. We used a random subsample of 500 probes and a leave-one-out cross- validation with AIC as the validating criterion. Then, we filtered probes that displayed a differential expression (significantly different from 0 after centering) in both lines. The following analysis is carried out on the mean expression of both lines. To detect a group of probes with similar variations, we chose to cluster the expression curves with kmlShape [22]. Some clusters displaying similar patterns were then manually pooled after visual inspection. The results of differential expression and clustered spline profiles are presented in S1 Table.

In the second step, a differential analysis was carried out to identify the DEG between d2 and d13 using the packages limma [23] and variancePartition [24], the second one providing an extension of the first one for linear mixed modeling. According to previous results and testing, we chose a model including time (i.e., duration of HS), line as fixed effect, and replicate and pig as random effects. After fitting this model on each probe, we filtered the results by keeping only probes with a false discovery rate adjusted p-value (FDR lower than 5%). All results are presented in S2 Table.

A functional enrichment analysis was performed on each DEG list for each cluster of the step one analysis with the package gProfiler2 [25]. We used the Gene Ontology (GO) [26], Kyoto Encyclopedia of Genes and Genomes (KEGG) [27] and Reactome [28] databases in our analysis. All the 32,924 probes corresponding to 10,087 unique Ensembl IDs that passed the filtering step during preprocessing were used as background for this enrichment analysis. To generate the plot that illustrates the enrichment results, it was first necessary to filter the enrichment terms, eliminating those that included fewer than 100 genes to exclude overly generic terms. Thereafter, the enrichment terms that exhibited overlap with a single gene within the query were eliminated. Finally, only the ten enrichment terms with the highest gene ratio (defined as the proportion of genes in the cluster present in the enrichment term relative to the total number of genes in the cluster) are presented. A more complete list of the identified enrichment for each cluster (step 1) is provided in S3 Table.

Finally, the DEGs from step 1 (d0 vs d2, STHA) and step 2 (d2 vs d13, LDHA) were compared using Qiagen Ingenuity Pathway Analysis software. This was done in order to highlight the canonical pathways that were differentially (FDR < 0.05) affected depending on the time of heat exposure, namely STHA vs LTHA. Ten canonical pathways were selected to be presented in Table 1, illustrated with a z-score to infer an activation state, positive values for increased or negative values for decreased of the predicted function. A z-score is conventionally considered significant if above an absolute value of 2. Results are presented in S4 Table.

**Table 1.**
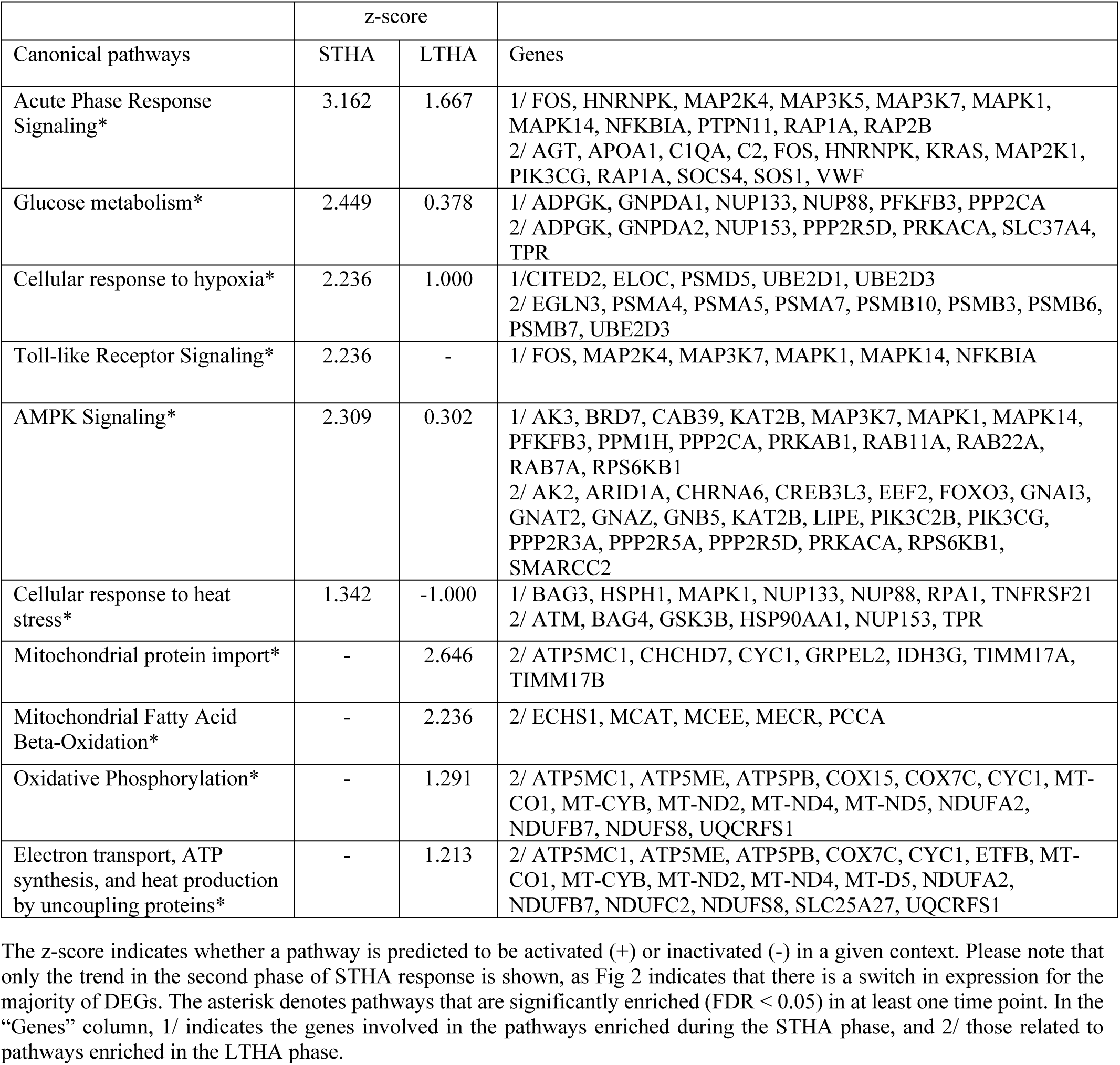
Selection of 10 canonical IPA pathways, comparing acclimation responses during STHA and LTHA periods.

## Results

### Thermoregulatory Responses

One of the most commonly used parameters to assess the thermal balance, as well as one of the simplest ones to determine, is rectal temperature (RT). As presented in Fig 1, irrespective of the line effect, RT began to rise during the first 2 days of exposure at 30°C and then slowly decreased over time. While both RFI lines displayed a similar general pattern of RT, a significant difference was observed between the two lines for day 2, day 6, and day 10 (p<0.05). Peak RT was reached on day 1 in LRFI pigs, whereas animals from the HRFI line reached peak RT 24 hours later. Furthermore, average RT was significantly higher in HRFI than in LRFI pigs during the whole experiment (39.8 vs 39.6°C, p=0.016) and heat exposure (39.9 vs 39.7, p=0.019). These differences between lines are not discussed in the present paper, which focuses mainly on generic acclimation responses to thermal HS.

**Fig 1.**
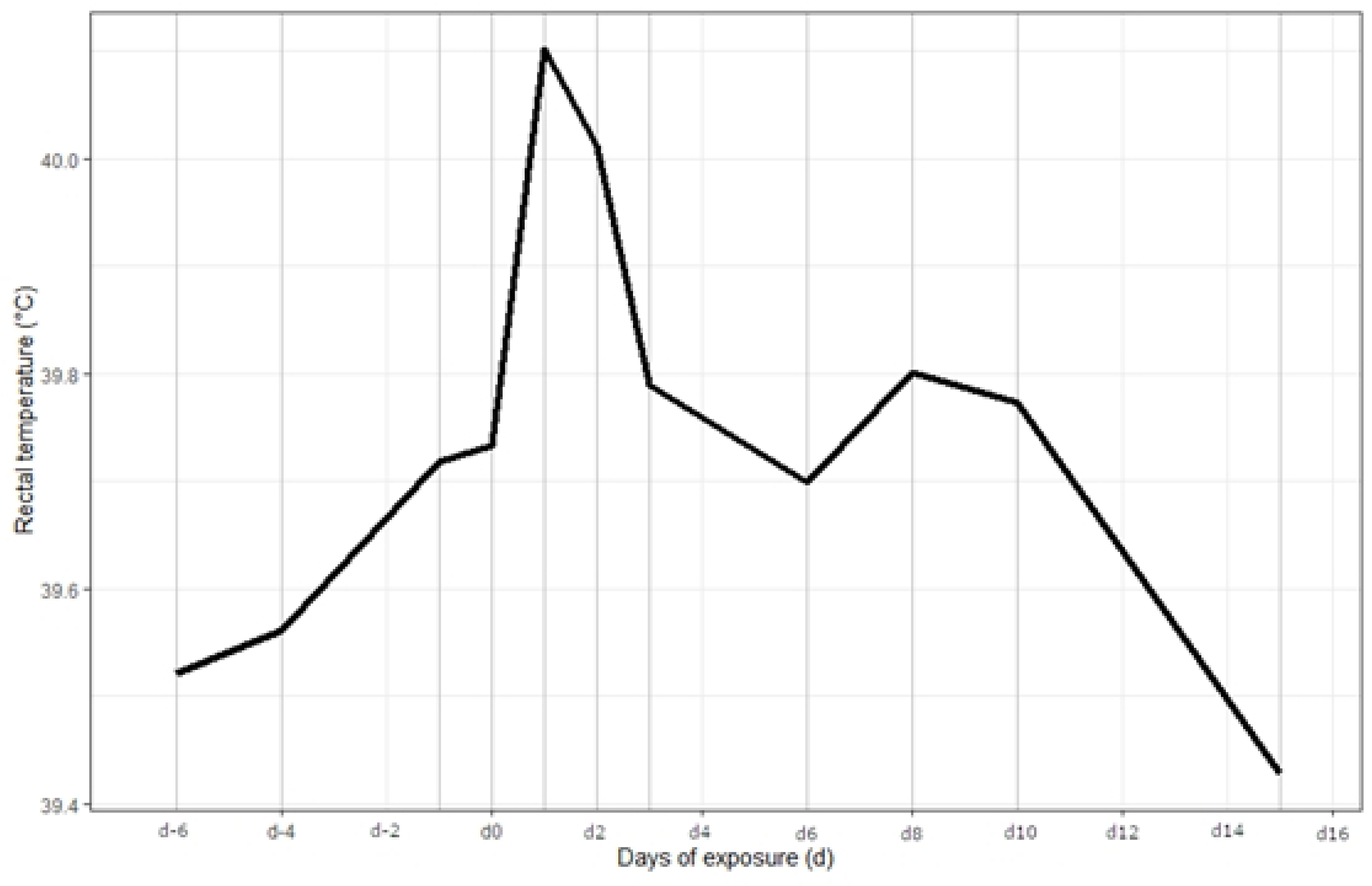
Effect of duration exposure at 30 °C on rectal temperature (n = 36 pigs). Day 0 = day of transition temperature between 24 and 30°C.

### Short-term whole-blood gene expression responses

The first step was to consider the kinetics of gene expression measured in blood cells during the first 48 hours of exposure to 30°C. We selected only the probes (n=935) that were differentially expressed in both pig lines, assuming that the associated genes (n=525) represented a shared, generic response across both lines.

A clustering method was applied to each DEG to identify groups of probes sharing a similar pattern of expression, i.e., GAM (Generalized Additive Model) modeled expression with a resembling shape. While the algorithm performed remarkably well given the number of expression patterns, a manual pooling of some clusters sharing a similar shape was performed before further analysis (Fig 2).

**Fig 2.**
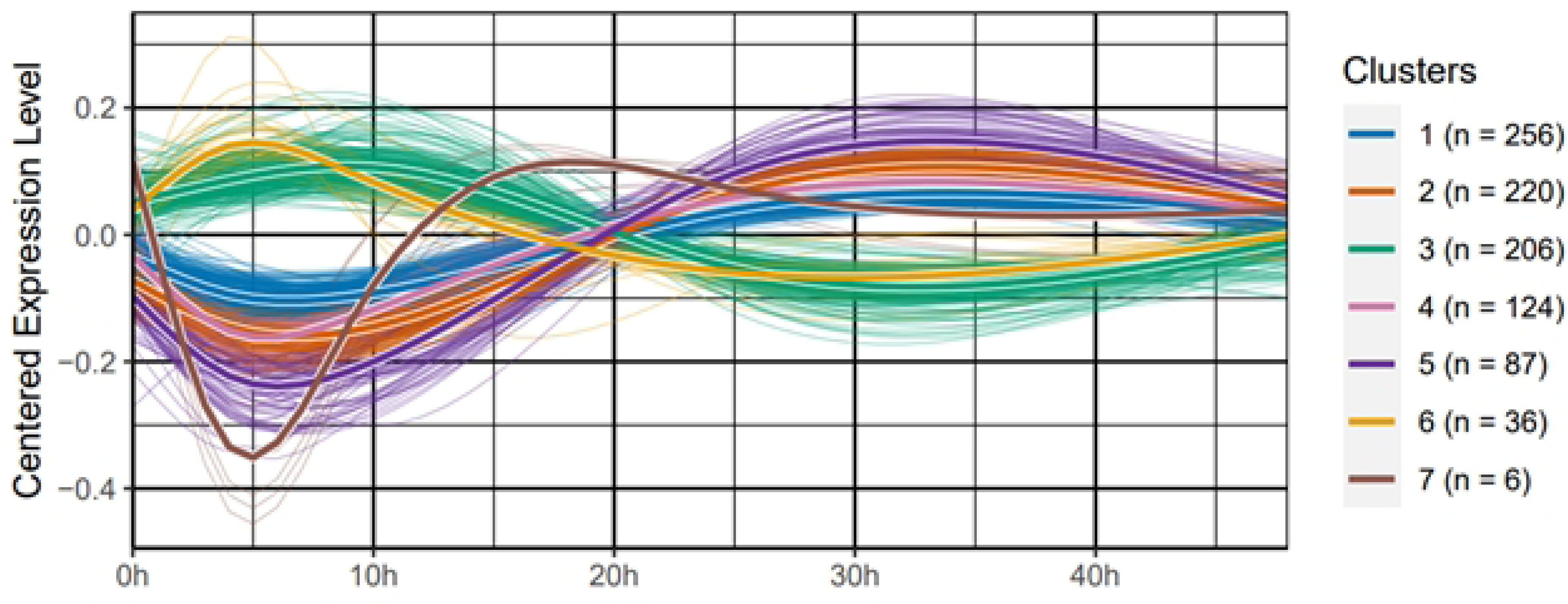
Patterns of expression during STHA of the 935 differentially expressed probes for both RFI lines (n = 12 pigs). The kinetics of gene expression are grouped into clusters according to their trajectories during STHA periods using the kmlshape package. Bold lines correspond to the average kinetics for each cluster. The number of unique differentially expressed genes per cluster is given in parentheses.

From the clustering analysis, a total of seven groups of gene expression were identified (Fig 2). Interestingly, our results reveal two predominant expression patterns with a switch occurring around 20 hours. A total of 687 genes of the clusters 1, 2, 4, and 5 were down-regulated during the first 10 hours, followed by an up-regulation until the 30th hour of exposure. Symmetrically, the cluster 3 exhibited 206 genes that were initially up-regulated, then progressively repressed. Genes aggregated in clusters 6 (36 genes) and 7 (6 genes) showed specific dynamics characterized by a faster modulation of their expression. Expression profiles are slightly different, with a more rapid increase and decrease for cluster 6 and the opposite for cluster 7. The cluster 7 consists of only two genes, i.e., Interleukin 1β and the immediate-early response gene C-Fos.

### Functional Enrichment during STHA Phase

Significant terms and pathways enriched from genes of the clusters are mainly linked to the immune system (clusters 1 and 2) and cell organization (clusters 4 and 5, Fig 3). Enrichment for clusters 1 and 2 is also related to the activation of the innate immune system through toll-like receptor signaling pathways. No enrichment terms were found for the cluster 3. However, it is interesting to observe that up-regulated genes between 0 and 20 h, i.e. cluster 3 and 6, there is the SMAD signaling pathway enriched term for cluster 6 (Fig 3) with SMAD2 (transducer and transcriptional modulator) and SP1 (transcription factor) genes, while the gene BMP4 (a secreted ligand of the TGF-beta (transforming growth factor-beta) superfamily), which is activated by SP1 and regulates SMAD pathway, belongs to cluster 3.

**Fig. 3.**
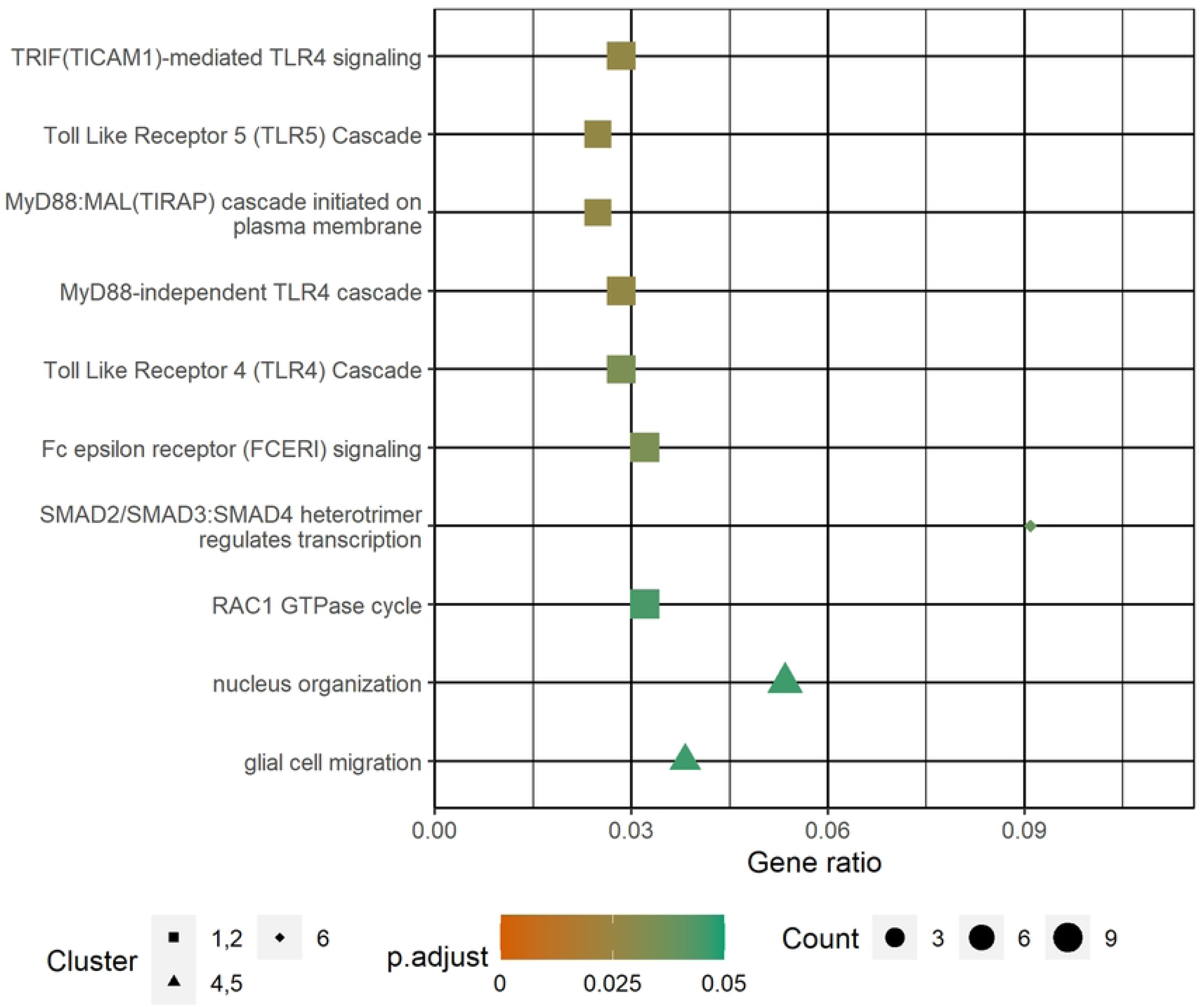
Enrichment analysis of the DEG aggregated in clusters 1, 2, 4, and 5. Point shape corresponds to the clusters presented in Fig 2. Point size translates to the number of genes in the cluster that intersect with the enrichment term.

### Long-term Whole-Blood Gene Expression Responses

From d2 to d13, 2,087 probes (FDR < 5%) were found to be differentially expressed, representing a total of 985 unique genes. A total of 825 probes (430 unique genes) exhibited decreased expression, and 1,262 probes (566 unique genes) showed increased expression. A small subset of genes (∼1%) is represented by both up- and down-regulated probes, which may reflect alternative transcript variants or hybridization artifacts. This explains the discrepancy between the total number of regulated genes (985) and the sum of the number of genes classified as either up-regulated (430) or down-regulated (566), which totals 996.

### Comparative Analysis of Molecular Pathways Involved During the HS Response

The STHA and LTHA DEG lists were subjected to functional exploration with the aim of identifying regulated molecular pathways that significantly differ between STHA and LTHA phases. A comparative process was carried out using Ingenuity Pathways Analysis, and the results are summarized in S4 Table. Table 1 shows a selection of 10 canonical pathways. Pathways that were significantly enriched for at least one condition, i.e., during the STHA phases (d0 to d2) and LTHA (d2 to d13), and/or with a z-score indicating an up- or down-regulated state of the pathway were retained in Table 1 for further discussion. Many signaling pathways have a negative and then a positive z-score for the STHA phases, illustrating an activation state just before the LTHA phase. During the LTHA period, the z-score of these pathways is reduced (lower positive z-score below 2) or even negative, indicating an inactivated state.

The z-score indicates whether a pathway is predicted to be activated (+) or inactivated (-) in a given context. Please note that only the trend in the second phase of STHA response is shown, as Fig 2 indicates that there is a switch in expression for the majority of DEGs. The asterisk denotes pathways that are significantly enriched (FDR < 0.05) in at least one time point. In the “Genes” column, 1/ indicates the genes involved in the pathways enriched during the STHA phase, and 2/ those related to pathways enriched in the LTHA phase.

Most of the selected molecular pathways of Table 1 are enriched at all kinetic points, except Toll-like receptor signaling, which is only active during the STHA period and mainly involves genes from clusters 1 and 2 (Fig 3). Other pathways underlie the state of active response to stress, e.g., hypoxia, acute phase, heat. Some others showed that some metabolic pathways are activated during the first two days of HS response, e.g., glucose and lipid metabolism. Most of these pathways involved AMPK signaling, several MAP kinases (e.g., with *MAP3K7, MAPK1,* and *MAPK14* genes), and PI3 kinase (e.g., *PIK3C2B* gene) as cellular checkpoints. Most of these signaling pathways are involved in the regulation of metabolic pathways. The metabolism of glucose is mainly up-regulated in late STHA, and the metabolism of lipids is up-regulated in late STHA and down-regulated in LTHA, with e.g., the lipase E, hormone-sensitive *LIPE* gene, and the perilipin *PLIN1* gene.

Another interesting pathway at LTHA is the oxidative phosphorylation in the mitochondria (FDR < 0.05). The z-score is 1.291 and, as shown in Fig 4, complexes I, III and IV are mainly inactivated, whereas Complex V is activated. More specifically, within the complexes I and III, genes encoding for NADH dehydrogenase subunits are up-regulated (*NDUFA2, NDUFB7, NDUFS8*), whereas the *MT-ND2* (Mitochondrial NADH dehydrogenase subunit 2) gene is down-regulated. Within the Complex III, the *UQCRFS1* gene encoding for the Cytochrome C1 is up-regulated and the *MT-CB* (Mitochondrial Cytochrome b) gene is down-regulated. Finally, the *COX1* (Cytochrome c Oxidase Subunit I) gene is down-regulated, while the *COX7* gene is up-regulated, as is the gene encoding Cytochrome C, *CYTC*. The situation is much clearer for Complex V, where all the genes are either measured or predicted to be up-regulated.

**Fig 4.**
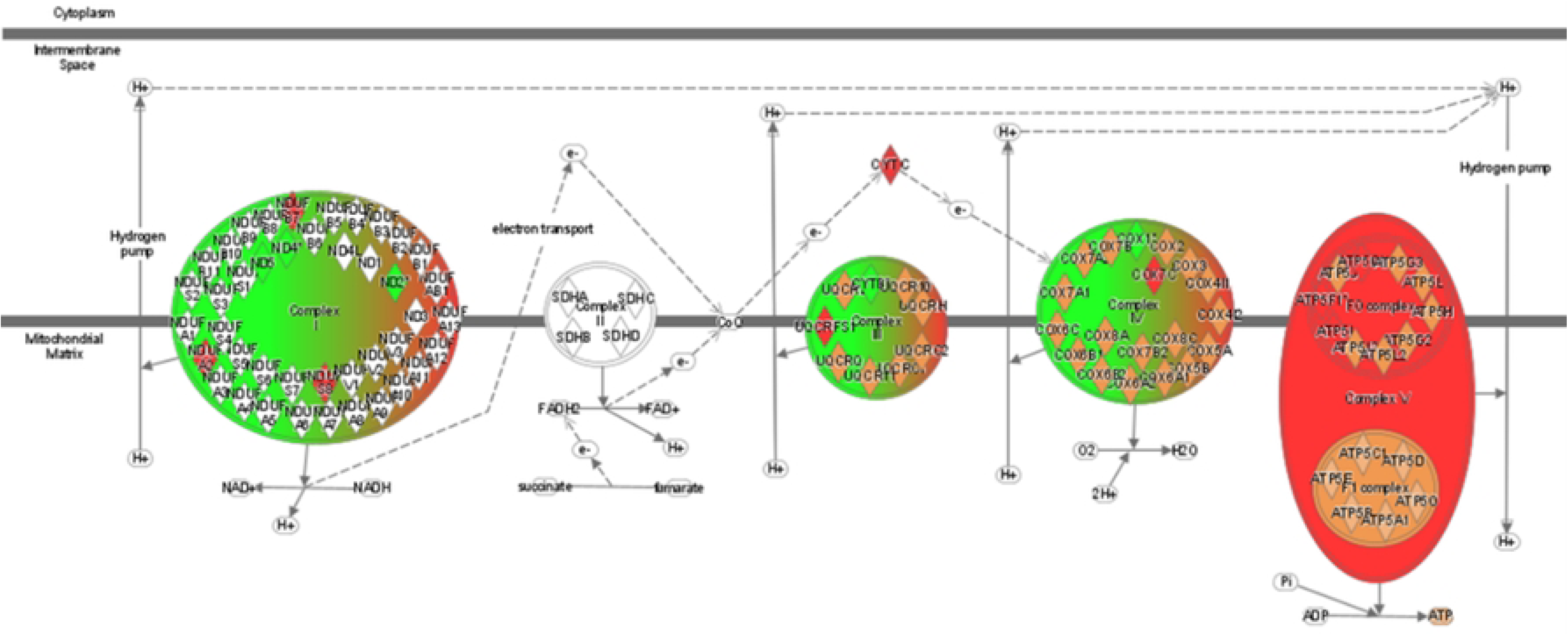
The oxidative phosphorylation pathway during the LTHA response. Green color indicates an inactivated state, red color an activated state according to the prediction by the z-score calculated under IPA.

## Discussion

Exposure to high temperatures results in various physiological and metabolic responses affected by the nature and severity of HS. As evidenced by rectal temperature measurements, our current results demonstrated that pigs exposed to 30°C for 21 d developed biphasic acclimation process involving transient perturbed phases (STHA) followed by a long-lasting period during which acclimatory homeostasis is developed (LTHA). These different responses from the animal occur at distinct time points. Moreover, as shown by Campos et al. [13], feed intake was reduced in both lines when exposed to 30 °C. Therefore, patterns of changes in gene expression during heat exposure are related to both direct effects of HS and indirect effects related to feed restriction.

Although very few studies have been published on the subject in pigs or other livestock species, these acclimation responses likely vary significantly between individuals [13]. It is therefore necessary to study the dynamics of these mechanisms within the same animal. This requires serial sampling over time on the same individual. However, except for easily accessible tissues such as the blood, serial sampling of other tissues, e.g., muscle, cannot be readily obtained. This presents a significant challenge for transcription-based investigations of complex and dynamic traits such as thermoregulation. To achieve this, we assume that a comprehensive view of the molecular mechanisms involved in the animal’s responses during HA can be obtained based on its whole blood transcriptome. According to a recent study on humans [29], the whole blood transcriptome could provide a significant prediction of tissue-specific gene expression for approximately 59% of genes on average across 32 tissues, with a maximum of 81% of the genes in the skeletal muscle. Similar findings have also been reported in pigs, leading us to assume that gene expression profiles in blood largely reflect those in other tissues, mainly the ones involved in immune and inflammatory response, to a lesser extent with intestine tissues, liver, and kidney [30]. In a former study, we performed a multi- tissue analysis of pig before and after a short-term heat stress [31]. We observed that most shared regulated genes were found in both muscle and adipose tissue, and to a lesser extent, shared regulated transcripts were also found in muscle and adipose tissue in conjunction with blood [31]. To go further, predictive models of tissue gene expression based on blood gene expression were developed and allowed to predict up to 60 % on average for 32 tissues in humans [29]. Based on these results, we hypothesize that the heat adaptation mechanisms elucidated from the blood transcriptome may be partially extrapolated to other tissues.

Our study delineates for the first time the dynamic gene expression measured in blood cells during different phases of HA in swine. Our results show that expression changes occur rapidly after the elevation of temperature and that the first effects of HS were predominantly inhibitory, as more than three quarters of the DE genes (405/525; clusters 1, 2, 4, 5, and 7) were suppressed during the first phase of STHA. This result is consistent with findings obtained in humans [32]. The transient gene down- regulation during the first 10 hours is followed by an increased expression of most genes. That suggests that shortly after the start of heat exposure, cells reprogram their molecular functions to activate early defense mechanisms. These cellular stress responses entail the activation of various signaling pathways, which facilitate the management of cellular damage and the initiation of repair processes. The Ingenuity Pathway Analysis identified MapK as one of the top networks signaling hubs for the cellular response to HS and hypoxia during the second STHA phases (i.e., 10 h after the onset of the thermal challenge). Various mitogen-activated protein kinases are up-regulated several hours after the onset of the thermal challenge and probably collectively contribute to the cellular and systemic responses to HS by regulating inflammation, stress signaling, and cellular adaptation mechanisms [33]. The IPA revealed that the canonical pathway related to the cellular response to HS was predicted to be significantly enriched but was moderately activated (Z-score =1.34) after 10-h exposure to 30°C (second phase of the STHA period). Surprisingly, only one HSP gene - *HSPH1* (Heat Shock Protein Family H member 1, also known as Hsp105) - was identified in this canonical pathway. Additionally, three differentially expressed genes encoding DNAJ HSPs (HSP 40 family members B12, C14, and C19) were found to be significantly up-regulated during the second phase of STHA. *HSPH1* and especially *DNAJ* homologs are considered one of the most highly conserved stress-response genes, as these proteins are required for stabilizing the interaction of HSP70 with unfolded proteins and enhancing the protective role of the HSPs [34]. In the present study, the *HSP70* gene tended (FDR p-value < 0.1) to be overexpressed, and probes belonging to the *HSP90* gene were either down-regulated or up-regulated during the STHA phases. However, these two proteins are known to be the most functionally relevant HSPs involved in the HSR [35], and their overexpression is classically reported in isolated peripheral blood mononuclear cells of pigs and other livestock species commonly incubated under extra physiological thermal conditions (>45°C) for several hours [36–38]. Similarly, the expression of *HSP70* was overexpressed in IPEC-J2 cells also incubated at 42°C for 2 hours [39]. In this study [39], the *HSP70* gene was also up-regulated in the colon tissue collected in piglets kept at 35°C for 21 days. This result is consistent with previous findings observed in gut or muscle tissues collected from HS growing pigs [3,40]. In fact, it can be assumed that the limited effects of STHA phases on the key HSPs gene expression may be attributed to reduced sensitivity of blood cells to HS compared to more metabolically active tissues such as muscle or digestive organs. Furthermore, we cannot exclude the possibility that the regulation of HSP expression in blood cells in response to HS varies significantly from one pig to another, and then the limited number of animals used in our study might have limited its statistical power.

In an attempt to improve heat dissipation, blood is redirected from the internal organs to the skin and the respiratory tract, where heat can be lost to the environment [41]. A reduction in blood flow to the viscera can result in a deficiency of oxygen delivery to these tissues, which may lead to a state of localized hypoxia in the gut and impair the integrity of the intestinal epithelium. This may result in an increased permeability of the gut, which is defined as a leaky gut. In pigs, this deterioration of the gut integrity is generally observed within the first 24 h of exposure to HS [3,42]. The heat-induced leaky gut could enhance the translocation of bacterial endotoxins and antigens into the bloodstream, which in turn can trigger systemic inflammation and immune responses. From these results, one can assume that the reprogramming of genes involved in the signaling pathways of the innate immune system (e.g. Toll-like receptor signaling or TLRs pathway) during the STHA phase may be indirectly related to the HS effects on gut integrity. In the present study, genes involved in the cellular stress response to hypoxia are up-regulated, confirming that cells must adapt to low oxygen levels during the second phase of the STHA period (between 20-48h after the stress induction). This reprogramming would be related to the above-mentioned thermoregulatory responses. In addition, hypoxia is known to activate Toll-like receptors and to increase the production of inflammatory cytokines, which can play a key role in initiating and modulating immune responses. Nevertheless, further investigation is required to substantiate this hypothesis, as the classic blood biomarkers of leaky gut (e.g., TNFα and lipopolysaccharide binding protein [43,44]) are not differentially expressed in the present study.

Our results indicated that genes related to glucose metabolism (*DLAT, DLD,* and *PFKFB3*) were up-regulated during the second part of the STHA phase. PFKFB3 (6-Phosphofructo-2-Kinase/Fructose-2,6-Bisphosphatase 3) is an enzyme that regulates the levels of fructose-2,6-bisphosphate (Fru-2,6-P2), a potent activator of phosphofructokinase-1 (PFK-1), the rate-limiting enzyme in glycolysis. Given the likelihood of transient cell hypoxia during the initial hour of HS exposure, it can be postulated that *PFKFB3* overexpression represents an adaptive mechanism to enhance glycolysis, thereby compensating for the diminished ATP production from oxidative phosphorylation [45]. In addition, *PFKFB3* expression is also reported in the case of oxidative stress. Under high levels of reactive oxygen species (ROS), PFKFB3 can be modified through S-glutathionylation, which decreases its enzymatic activity [46]. This results in a reduction of Fru-2,6-P2 levels, shifting glucose metabolism towards the pentose phosphate pathway (PPP). The PPP plays a pivotal role in the generation of NADPH, which is crucial for combating oxidative stress by maintaining the reduced form of glutathione. This, in turn, has a positive effect on ROS detoxification.

During LTHA (i.e., after 2 days of exposure to 30°C), the physiological responses of pigs to elevated temperatures are characterized by a reduction in hyperthermia and in reliance on evaporative heat loss. Additionally, a gradual increase in voluntary feed intake is observed [13]. From these results, it has been assumed that the gradual adjustment towards homeothermy was mainly related to a reduction in heat production, with animals generating less metabolic heat for the same amount of feed intake [13]. The observation of consistent results in different animal species indicates that modifications to the respiratory chain that occur during HA allow for a reduction in heat production [47]. According to the author, these modifications have to be connected to a reduced plasma concentration of thyroid hormones in acclimating animals. As detailed by Campos et al.,[13], T3 and T4 plasma levels significantly drop on the first day at 30°C, but showed a progressive increase in the following days. In addition, our findings suggest that LTHA exerts disparate effects on the enzyme complexes of the respiratory chain. The expression of the genes associated with Complex II remains unchanged during LTHA. Within each multi-subunit complex (I, III, and IV), gene expression of key enzymes of the electron transport chain—whether up-regulated or down-regulated—complicates the interpretation of our results. This also probably explains the low z-score associated with the “Oxidative Phosphorylation” pathway. In agreement with Chamberlin et al. [48], these results suggest that each reaction within oxidative phosphorylation may display different thermal sensitivities and that this variability could be amplified by the fact that individual animals may also have different responses. However, in accordance with Hafen et al. [49], all the genes of the component of complex V are either measured or predicted to be up-regulated during LTHA. The mitochondrial inner membrane-located ATP synthase catalyzes the terminal step of oxidative phosphorylation, producing energy-rich ATP from the electron transport-generated proton gradient [50]. Assuming that the activity of ATP synthase is directly related to the levels of gene expression encoding subunits of each complex, we hypothesize that the ATP synthase activity and respiratory capacity of mitochondria will increase during the LTHA period. This assumption is confirmed by results obtained in human skeletal muscles and reviewed by Marchant et al. [51]. Collectively, these findings highlight that mitochondria play a pivotal role in the long-term adaptation to HS.

## Conclusion

The present study aimed to explore the dynamics of the expression of the pig transcriptome during HS. Heat stress affected the expression of various genes at both short-term and long-term. A coordinated response seems to take place after one day of exposure to a hyperthermic environment; a shift of expression was observed. Comparative pathway analysis indicated distinct biological responses between short- and long-term heat acclimation. These findings suggest a temporal regulation of stress and metabolic pathways, with STHA involving acute heat shock responses and metabolic adjustments to support homeothermy, and LTHA reflecting more sustained bioenergetic reprogramming. This study contributes to a better understanding of the timeline of adaptation mechanisms of pigs to HS. Further studies with more frequent sampling and a focus on the first two days of HS would be of particular interest to better map the expression dynamics of the genes involved in high-temperature adaptation.

## Supporting information

**S1 Fig: Experimental design.**

**S1 Table**: **Table of the DEGs identified during short-term HS in the spline analysis.** DEG are clustered according to their profile.

**S2 Table: Table of the DEGs identified between d2 and d13.** Adjusted p-value cutoff was set to FDR < 0.05.

**S3 Table: Table of enrichments identified in each cluster**. Pathways were enriched with GO, KEGG, and Reactome databases. Cutoff for adjusted p-value was fixed at 0.05. A background corresponding to the annotated genes was used to correct multiple test p-values.

**S4 Table: Table of the Ingenuity Canonical Pathways identified in both STHA and LTHA.** Adjusted p-values and z-scores are given. A cutoff for adjusted p-value was fixed at 0.05, or for the absolute value of the z-score was fixed above 2 in at least one STHA or LTHA condition.

## Acknowledgments

The authors gratefully acknowledge Loïc Gaillard, Francis LeGouevec, Alain Chauvin, Régis Janvier, Serge Dubois, and the GENESI staff for animal care and technical assistance, Aline Foury, Laure Gress, Anne Pasquier, and Christine Tréfeu for the laboratory analysis and Jerome Lecardonnel of the @Bridge platform for performing the transcriptome analysis.

